# Human Olfactory Perception is Sensitive to Temporal Information within a Single Sniff

**DOI:** 10.1101/2020.08.29.271254

**Authors:** Yuli Wu, Kepu Chen, Kai Zhao, Wen Zhou

## Abstract

A sniff in humans typically lasts 1-2 seconds and is considered to produce a “snapshot” of the chemical environment that also sets the temporal resolution of olfactory perception. To examine whether the temporal order of events within a sniff influences the perceptual “snapshot”, we devised an apparatus that enabled us to phase-lock odor delivery to sniff onset and precisely manipulate onset asynchronies of odorants in humans. Psychophysical testing showed that participants were able to tell apart two odorants presented in the same or different order when the onset asynchrony was as low as 40 milliseconds. The performance improved with longer onset asynchronies and was not based on the molar ratio difference of the two odorants. Meanwhile, they were consistently at chance in reporting which odorant arrived first. These results provide behavioral evidence that human olfaction is sensitive to temporal patterns within a single sniff and indicate that timing of odor-evoked responses in relation to the sniff contributes to the perceived odor quality.

## Introduction

An important computational challenge for human brain lies in processing the rapidly changed exogenous sensory information. Most sensory systems, including visual and auditory, have a fine temporal resolution in sensory processing, which allows us to quickly react to the external world (Kanabus, Szelag, Rojek, & Poppel, 2002). While, human olfactory is thought as coarse in this progress with a discontinuously sampling of information, namely sniffing that typically lasting 1-2 seconds. This has been shown in previous studies that we have difficulty in distinguishing the sequence of two successively presented odor, which may because of the suppression between odors (Getchell, Margolis, & Getchell, 1984; Laing, Eddy, Francis, & Stephens, 1994). Thus, human olfaction was also regarded as a change blinded state (Sela & Sobel, 2010). Although, a sniff sets a start of olfactory perception and supports related temporal information which may be extracted by human olfactory system, few clear evidence shows that it can substantially changes the odor perception (Perl, Nahum, Belelovsky, & Haddad, 2020).

However, more and more animal models revealed a fine temporal sensitivity and the involvement of temporal details in the olfactory perception. In rodents, spiking latency, relative to the start of a sniff, stably encoded the identity of odorants in the olfactory receptor neuron and olfactory bulb (Bathellier, Buhl, Accolla, & Carleton, 2008; Bolding & Franks, 2018; Cury & Uchida, 2010; Iwata, Kiyonari, & Imai, 2017; Shusterman, Smear, Koulakov, & Rinberg, 2011). Furthermore, relative temporal information from olfactory bulb can be read out by the olfactory cortical neurons (Haddad et al., 2013). Correspondingly, manipulating the stimulator‘s onset phase in a sniff within tens of milliseconds can get different behavior and neural/neuronal outputs in rodent (Smear, Resulaj, Zhang, Bozza, & Rinberg, 2013; Smear, Shusterman, O’Connor, Bozza, & Rinberg, 2011), which also indicates that phase is a non-negligible factor in determining the role of temporal information and its sensitivity in olfactory perception.

Therefore, in this study, we designed an apparatus that enabled us to deliver odor phase-locked to sniff onset and precisely manipulate onset asynchrony of two odorants to examine whether the temporal order of events within a sniff influences odor quality perception. What’s more, we further set out to assess the sensitivity of human olfactory system to this temporal information in olfactory perception.

## Results

### The apparatus for odor presentation is sensitive in temporal control

In order to deliver odors precisely, asynchronously and locked to sniff, we designed an apparatus (see details in Fig. 1A and method) and verified its availability and stability in time control. This apparatus controls the odor onset asynchrony by manipulating tubes length differences (ΔL) and actively sniffing. We measured vapor phase odor concentration to test the onset asynchrony (SOA) between the odors with two calibrated miniature photo-ionization detectors (miniPID 200B, Aurora Scientific, Canada) (Fig. 1A). Flow rate of the pump was set at 100 ml/s, based on our pilot measurement of nasal flow rates when sniffing through the apparatus, whereas ΔL varied from 0 to 40cm. As a result, odor concentration changed differently in two tubes after the pump turned on (Fig. 1B). We tested the SOA in a series of tube length differences from 0cm to 40cm for 32 times per length (Fig. 1B). Then we fitted these SOAs (ms) with ΔLs (cm) and got a liner relationship: SOA =3.9*ΔL +2.1 (Fig. 1C). Even when ΔL=5cm, the onset time in the short tube is early than the longer one for about 20ms (t_31_=4.909, p<0.001). When ΔL increased to 40cm, they have an SOA around 158 ± 9.7 (Mean ± SEM, ms). Therefore, by using of two mini PIDs, we confirmed that our apparatus is well in controlling the SOA of two odors in a sniff and got the liner relationship between ΔL and SOA.

**Figure 1:**
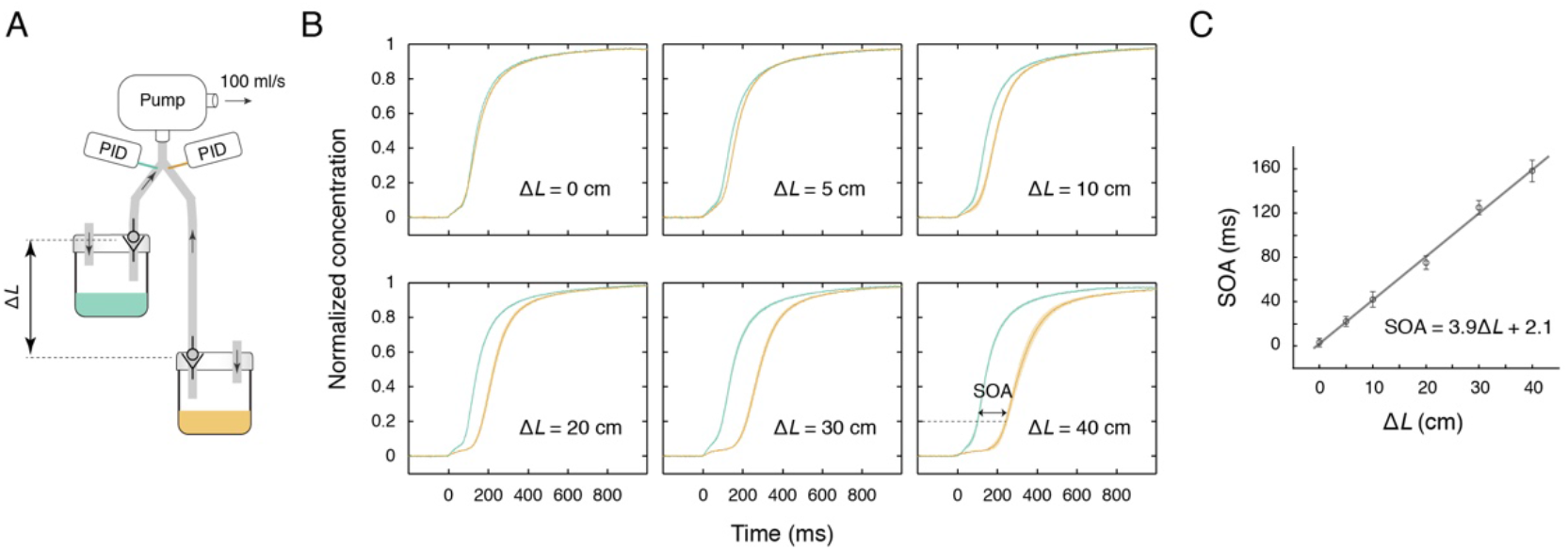
(A) Schematic illustration of the apparatus and SOA measurement. Vapor phase odor concentrations in the two tubings were measured close to the Y joint by mini PIDs. (B) Comparisons of the odor concentration time courses in the two tubings at different Δ*L* values. SOA was calculated as the time interval between when the odor in the short tubing reached 20% of its max concentration and when that in the long tubing reached 20% of its max concentration. Time 0 marks when the vacuum pump was turned on. Shaded areas: SEMs. (C) SOA increased linearly with Δ*L*. Error bars: SEMs.

### Phase difference can affect odor perception in a single sniff

Then, in experiment 1, we want to know if the temporal order of two odorants can affect olfactory perception within a single sniff for human beings. We selected two odorants, which have similar intensity (p>0.8) and valence (p>0.7) rating, and trigeminal property (see supporting information), including 10% v/v Isobutyl Phenylacetate (IP) and 2% v/v Pentyl Valerate (PV) dissolved in propylene glycol and stored separately in two bottles of the apparatus. The order was manipulated by which odor bottle was connected to the short tube. ΔL was set as 30 cm, which was estimated to produce an SOA of ~120 ms. In each trial, the two odorants were presented twice to the same nostril (the other nostril was pinched shut), in the same or different orders, and participants indicated whether the odors smelled the same or different (Fig. 2A). The residual odorants in the apparatus were cleared after each sniff. Participants need to finish 8 trials, after a practice procedure with feedback. Data from 24 participants showed that they had a significant higher accuracy than chance level (chance = 0.5) in the judgement of whether these two smells were same or not (Fig. 2B; mean accuracy = 0.688, p < 0.001, t_23_ = 6.044, Cohen’s d = 1.234; BF_10_ = 5449, error = 3.087e-8%). In other words, although participants sniffed the same odor mixture of IP and PV but in different temporal order, they felt different, which indicates the involvement of phase details in odor encoding.

**Figure 2:**
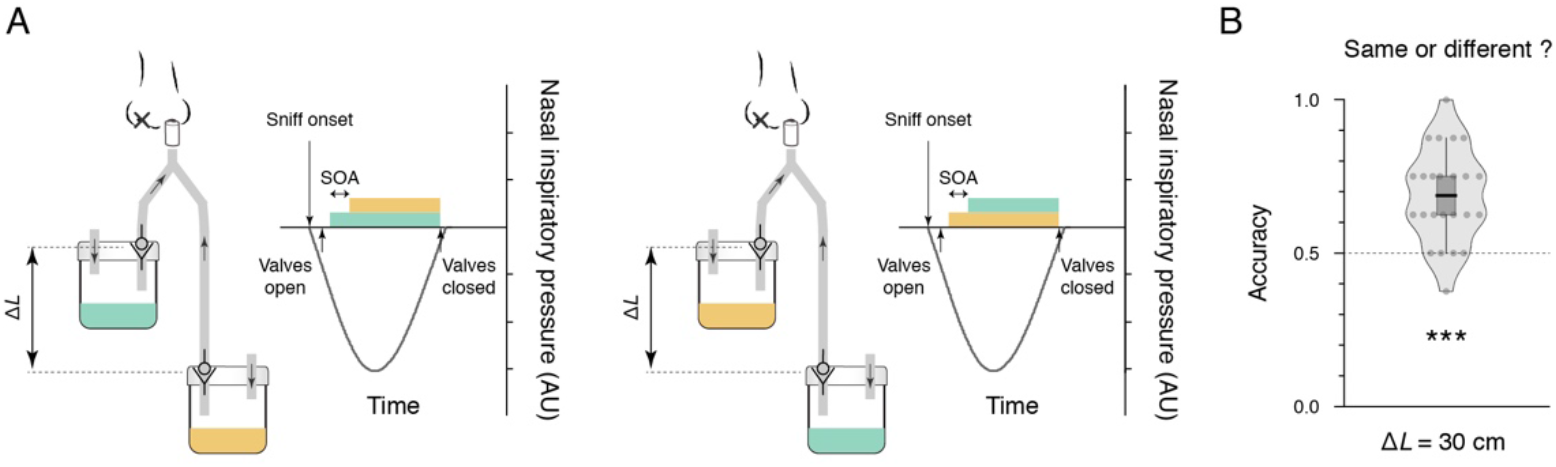
(A) Schematic illustration of an exemplar trial in the odor discrimination task. (B) Violin plots with included box plots of the odor discrimination accuracies of individual participants (gray cirles) in Exp. 2. The central black line denotes the mean, and the bottom and top edges of box indicate the 25^th^ and 75^th^ percentiles. Whiskers represent the data range expressed as the 75th percentile plus 1.5 times the difference between the 75th and the 25th percentile (maximum range) and the 25th percentile minus 1.5 times the difference between the 75th and the 25th percentile (minimum range). ***, p < 0.001.

### Human has a fine temporal sensitivity in a single sniff

Would human beings have a much better sensitivity to this temporal information? In experiment 2, we manipulated the ΔL, including 5 cm, 10 cm, 15 cm, 20 cm, 25 cm, 30 cm, and 40 cm, to control the SOA, in a similar distinguish task to experiment 1. To further characterize the temporal sensitivity of human olfactory perception, we recruited individuals with >=75% odor discrimination accuracies at ΔL = 30 cm to join in our formal experiment over 7 days. Apart from the discrimination task in which odor mixtures with same or different onset sequences, participants also had a sequence judgement task, in which they need to report which odorant was smelled first in one sniff. This task was set to investigate whether participants can clearly know the odor presentation sequence.

As shown in Fig. 3A, participants were able to tell apart two odorants presented in the same or different order in almost all SOA conditions. This capacity was impaired with the decreasing of SOA (F_6, 138_ =13.375, p<0.001) and dropped to chance level when the length difference is 5 cm (p>0.3; 5cm corresponded to an SOA of 20 ms). That is, when the ΔL is 10 cm, participants can still distinguish the odorants with different onset sequence over chance (Accuracy=0.64, t_23_ = 3.58, p<0.05, Cohen’s d=0.73, Bonferroni-Corrected; BF_10_ = 23, error = 2.792e-4%). The average airflow rate in tubes is closed to 50ml/s in each condition (all ps>0.38), indicating an SOA of 40 ms at ΔL = 10 cm. While the phase detail may work in a time scale within tens of millisecond to olfactory perception, participants consistently could not report which odor (IP or PV) was smelled first in all SOA conditions (Fig. 3C; all ps>0.15, Bonferroni-Corrected). Because of the similar respiration states in the discrimination task part and airflow testing part (Fig. S1, all p>0.05, FDR-Corrected), we also coarsely calculated the actual SOA for each participant at each ΔL based on per participant’s nasal flow rate (see method) and found a significant correlation between the actual SOAs and discrimination accuracies (Fig. 3B; r=0.459, p<0.001). These results indicate that human olfactory system may utilize the phase difference to encode odor identity with a sensitivity of tens of milliseconds.

**Figure 3:**
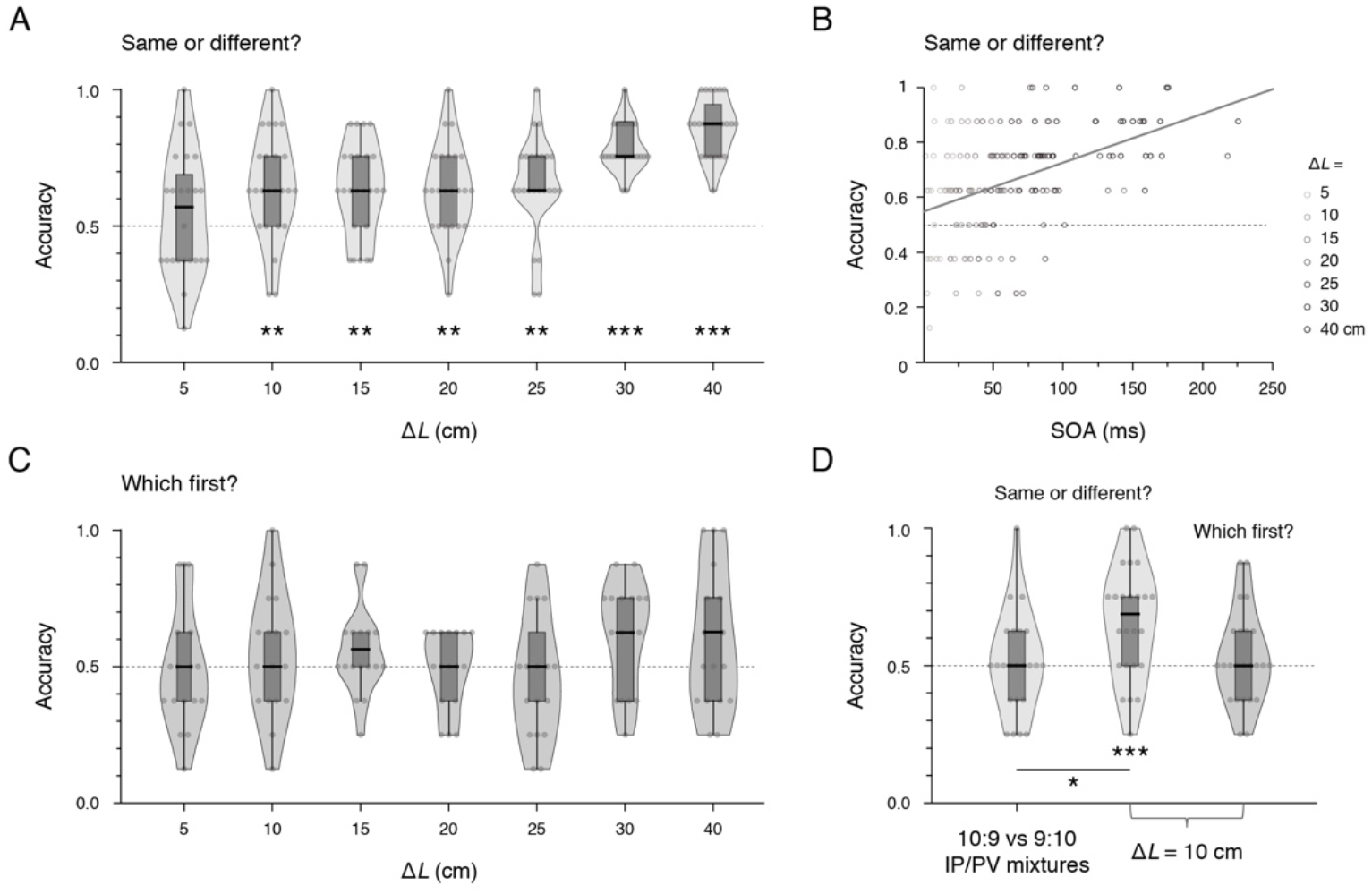
(A, C, D) Violin plots with included box plots showing participants’ accuracies in judging whether the odors were the same or different (A, D) and which odorant came first (C, D) in Exps. 3 (A, C) and 4 (D). (B) Odor discrimination accuracy correlated with calculated SOA in Exp. 3. *, p<0.05, **, p<0.01, ***, p < 0.001.

### This effect does not result from the molar ratio difference of two odors

Obviously, an early presented odor means more accumulated molar quantity, especially in the early stage of each sniff. Previous research claimed that a 420ms sniff is sufficient for odor identification (Laing, 1986). That is, the molar ration difference between the odor from via the short tube and the other via the long tube is no more than 10:9 in the first 420ms at ΔL = 10 cm based on the average SOA of 40 ms, which still bring different odor perception. To verified that the observed temporal sensitivity was not based on the molar ratio difference of IP and PV, we carried out Experiment 3 which has two olfactory conditions, one is same to the experiment 2 with ΔL = 10 which is called temporal difference, and the other condition is two kinds of odor mixtures of IP and PV in a volume ratio of 10:9 or 9:10 which is called molar ratio difference. Thus, participants needed to distinguish the odor mixtures in volume ratio of 10:9 with the opposite in the molar ratio difference condition. Participant were also chosen by the same criterion as Exp. 2. As a result, participants who performed significantly above chance at ΔL = 10 (Fig. 3D; t_23_=3.776, p<0.001, Cohen’s d=0.771; BF_10_ = 36, error = 1.983e-4%), nonetheless failed to differentiate between IP/PV mixtures in 10:9 and 9:10 volume ratios (Fig. 3D; t_23_=0.413, p=0.684, Cohen’s d=0.084; <u>BF_10_ = 0.232, error = 0.034%</u>). What’s more, accuracy in the temporal difference condition is significantly higher than the molar ratio difference condition (Fig. 3D; t23=2.427, p=0.012, Cohen’s d=0.495, one tailed; BF_−0_ = 4.703, error = 3.43e-5%, one tailed). Therefore, the SOA induced perceptual change may not due to the accumulated molar ratio difference.

## Discussion

An existed strategy often has its particular niche in perception. This was widely discussed in the processing of odor induced temporal pattern, leading to a temporal dynamic based coding strategy (Hopfield, 1995; Uchida, Poo, & Haddad, 2014). From mammals, amphibian to insects, temporal patterns are speculated for coding odor identity (Cury & Uchida, 2010; Junek, Kludt, Wolf, & Schild, 2010; Martelli, Carlson, & Emonet, 2013; Paoli et al., 2018; Su, Martelli, Emonet, & Carlson, 2011). What’s more, optogenetical methods were used to investigate the perception of temporal information in olfactory system, including mammals’ downstream neuronal activity and behavior output (Haddad et al., 2013; Smear et al., 2011). Nevertheless, few evidence directly gives point to the effect of odor temporal dynamic on mammal’s olfactory perception. In this study, by use of real odorants and phase locked odor presentation, we found human can distinguish the odor mixtures with different temporal dynamics although they did not know the sequence within it. This indicates that phase details of odorants presentation are resolved in olfactory processing. Moreover, participants also showed a fine sensitivity to temporal patterns in tens of milliseconds, suggesting a phase coding strategy in olfactory system.

A recent study also showed human can distinguish asynchronously presented odor mixtures with an SOA of 300 ms if they were informed with temporal feature (Perl et al., 2020). This is quite different from our result around 40 ms. We speculate the following reasons. Firstly, sniff is important for olfactory perception (Mainland & Sobel, 2006), and it induces sniff-coupled oscillations which benefit to robust phase coding of odors (Fukunaga, Berning, Kollo, Schmaltz, & Schaefer, 2012; Iwata et al., 2017) and cognitive function (Zelano et al., 2016). In our study, odor delivery was locked to sniff cycle and the onset phase is about 100ms after sniff based on our PID test, which may keep a higher olfactory sensitivity to temporal information. What’s more, our sniffing duration is longer which is likely to increase the discrimination accuracy (Laing, 1986; Rinberg, Koulakov, & Gelperin, 2006; Sobel, Khan, Hartley, Sullivan, & Gabrieli, 2000). At last, although we excluded that molar ratio difference is not the determining factor for successfully discriminating these odor pairs, we acknowledge that there is no absolute reason to rule out its contribution to higher sensitivity. A previous study also give us a worried that odorant with trigeminal property has a speed advantage in modulating sniff (Johnson, Mainland, & Sobel, 2003), we excluded the potential trigeminal factor by examining whether these two odors are same in trigeminal property (Hummel, 2000).

At the end, our study also put forward more thinking about whether asynchronous sensation is involved in more aspects. In the processing of odor mixture, animal models showed an integration of odor induced temporal profiles (Giraudet, Berthommier, & Chaput, 2002; P. Gupta, Albeanu, & Bhalla, 2015). Essentially, odorants have different crossing time in nasal mucosa which leads to an asynchronous of odor presentation similar to our experiment setting (Mozell & Jagodowicz, 1973). In our study, manipulating the odor temporal patterns results in a different olfactory perception, giving a supporting to holistic processing of temporal patterns in odor mixture perception. Apart from odor identity processing, asynchronous sensations from bilateral nostrils may also transmit spatial information, such as temporal difference based odor localization, which was found in animal studies that rats and sharks use the odor onset difference in two nostrils to get direction of the odor source (Gardiner & Atema, 2010; Rajan, Clement, & Bhalla, 2006). This leads to further researches to build a connection between time and space.

## Methods

### Participants

A total of 96 healthy nonsmokers took part in the study, 24 in Experiment 1 (12 females, mean age ± SD = 24.0 ± 3 yrs.), 24 in Experiment 2 (13 females, 23.2 ± 1.9 yrs.), 24 in Experiment 3 (13 females, 22.0 ± 1.7 yrs.) and 24 (12 females, 24.8 ± 3 yrs) in supplemental experiment. All participants reported to have a normal sense of smell, and no respiratory allergy or upper respiratory infection at the time of testing. Written informed consent and consent to publish was obtained from participants in accordance with ethical standards of the Declaration of Helsinki (1964). The study was approved by the Institutional Review Board at Institute of Psychology, Chinese Academy of Sciences.

### Olfactory stimuli

The olfactory stimuli consisted of isobutyl phenylacetate (Exps. 1-3, 10% v/v in propylene glycol), pentyl valerate (Exps.1-3, 2% v/v in propylene glycol), and their mixtures in 9:10 and 10:9 volume ratios (Exp.3), respectively, i.e., one containing 4.7% v/v isobutyl phenylacetate and 1.06% v/v pentyl valerate, and the other containing 5.4% v/v isobutyl phenylacetate and 0.94% v/v pentyl valerate. The tracer odor used in PID test of our apparatus was 1% v/v Isoamyl Acetate which was suitably used in testing temporal feature (P. Gupta et al., 2015; Priyanka Gupta, Albeanu, & Bhalla, 2016). They were presented in identical 200 ml glass bottles. Each bottle contained 10 ml of clear liquid.

### Apparatus

The apparatus (Fig. 1A & Fig. 2A) contained two tubings (ID = 4.5 mm), one 10 cm and the other 10 + ΔL cm in length. They were connected to a mini vacuum pump (in apparatus verification and for odor residue removal in Exps. 1-3) or a Teflon nosepiece (during odor presentations in Exps. 1-3) via a Y structure. The free ends of the two tubings were each fitted with a one-way valve and a push-to-connect tube fitting that allowed for easy connection and disconnection to an odor bottle. Odors could not emanate into the tubings unless the check valves were opened by a negative pressure exerted by the pump or by sniffing through the nosepiece.

### Procedure

In experiment 1, 2, and 3, participants need to finish an odor discrimination task with odor onset asynchrony (SOA, temporal difference) induced by tube length difference (ΔL, 30cm for Exp.1; 40, 30, 25, 20, 15, 10, and 5 cm for Exp.2; 10cm for Exp.3) or volume ratio difference (10:9 for experiment 3) with 8 trials of each. In each trial, participants need to sniff through the apparatus using one nostril twice (the other nostril was pinched shut) with a 2-4 seconds interval and make a forced choice whether the smells are same or not. For the convenience of odor presentation, we have two theoretical identical apparatus for these two successively sniffs. After each trial, there was a break for more than 1 minutes, when experimenter need to clear up the apparatus by pump during the break. Before each formal experiment, participants have practice trails with feedback.

Besides, in experiment 2 and 3, participants’ respiration state was monitored by respiration belt (Biopic MP150, Goleta, CA) during the formal experiment of odor discrimination task. After the odor discrimination task, the input terminals were connected with NV1 Rhinospirometer to test the nasal flow rate during sniff, while participants were asked to sniff as in former experiment and their respiration state was also monitored by respiration belt. Furthermore, they also have an odor sequence task under each SOA, in which they need to smell through the apparatus once and oral report which odor is smelled first with 8 trials in total, after the odor discrimination task. Before the odor sequence task, participant need to smell IP and PV separately to familiar with these two odors. All length difference conditions were finished in different days with no need in successive days.

### Analyses

The discrimination accuracy in each condition were compared with chance = 0.5 in a series one sample t-tests with Bonferroni-Corrected, two tailed. The respiration states between task part and airflow testing part were compared by paired samples t-test with FDR-Corrected.

We calculate the actual SOA by using of the nasal flow rate of each participant tested after the odor discrimination task. NV1 Rhinospirometer can get an inspiration capacity in a lasting time. We divided the inspiration capacity by the sniff time to get the airflow rate in each tube. The tube’s cross-sectional area is S, so we can get the speed of the odor in each tube roughly: Speed = Velocity / S. Then the length of each tube divided by the speed in each tube is the onset time of each odorant after sniff. So that, we get the onset asynchrony time of each subject (SOA = L_*L*_ / Speed_*L*_ − L_*S*_ / Speed_*S*_).

### PID measurements

Two calibrated miniature photo-ionization detectors (mini PIDs, Aurora Scientific, Canada) were used to measure the time onset asynchrony at the convergent part (Fig. 1A). The probe needles of each mini PID were respectively inserted into the converge part of these two tubings to test changes of vapor phase odor concentrations after sniffing. Sniff was simulated by a suction pump with an airflow rate of 100 ml/s, which bring an airflow rate in 50 ml/s for each tube. Therefore, we tested the SOA of odors in various tubing length differences, including 0, 5, 10, 20, 30, and 40 cm, each for 32 trials. The output voltage was corrected by this equation: 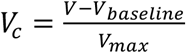, where *V*_*baseline*_ is defined by the mean voltage from −200 ms to −50 ms, and *V*_*max*_ is defined by the maximum from 0 ms to 2000 ms after bump opened. Onset time was defined by the time when *V*_*c*_ = 0.2 after bump opened.

## Supporting information

Supplemental experiment and figure

## ACKNOWLEDGMENTS

We thank Yan Zhu for supporting mini PID. This work was supported by the Key Research Program of Frontier Sciences (QYZDB-SSW-SMC055) and the Strategic Priority Research Program (XDBS01010200) of the Chinese Academy of Sciences, the National Natural Science Foundation of China (31830037) and Beijing Municipal Science and Technology Commission.

