## Supplemental experiment and figure for "Human Olfactory Perception is Sensitive to Temporal Information within a Single Sniff"

To estimate whether odor IP and PV have similar trigeminal property or not, participants performed two tasks based on odor lateralization. In the first task, participants were presented with IP and PV in bilateral nostrils separately. Then they need to make a judgment which nostril was presented with IP (or PV) in a total of 16 trials. Therefore, we can exam whether participants can lateralize these odors relied on the difference of trigeminal property. In the second task, we tested the trigeminal property directly. Participants were also presented IP and PV in bilateral nostrils separately, but they need to report which nostril feels more pungent. Then, we compared whether the nostril presented with IP was chosen as more pungent in an equal probability with that with PV. 24 participants performed these tasks. And we found they can neither lateralize these odors ($p>0.12$) nor have bias on choosing which is more pungent ($p>0.9$). Therefore, these two odorants should have no significant difference in trigeminal property.

*Supplemental Figure*

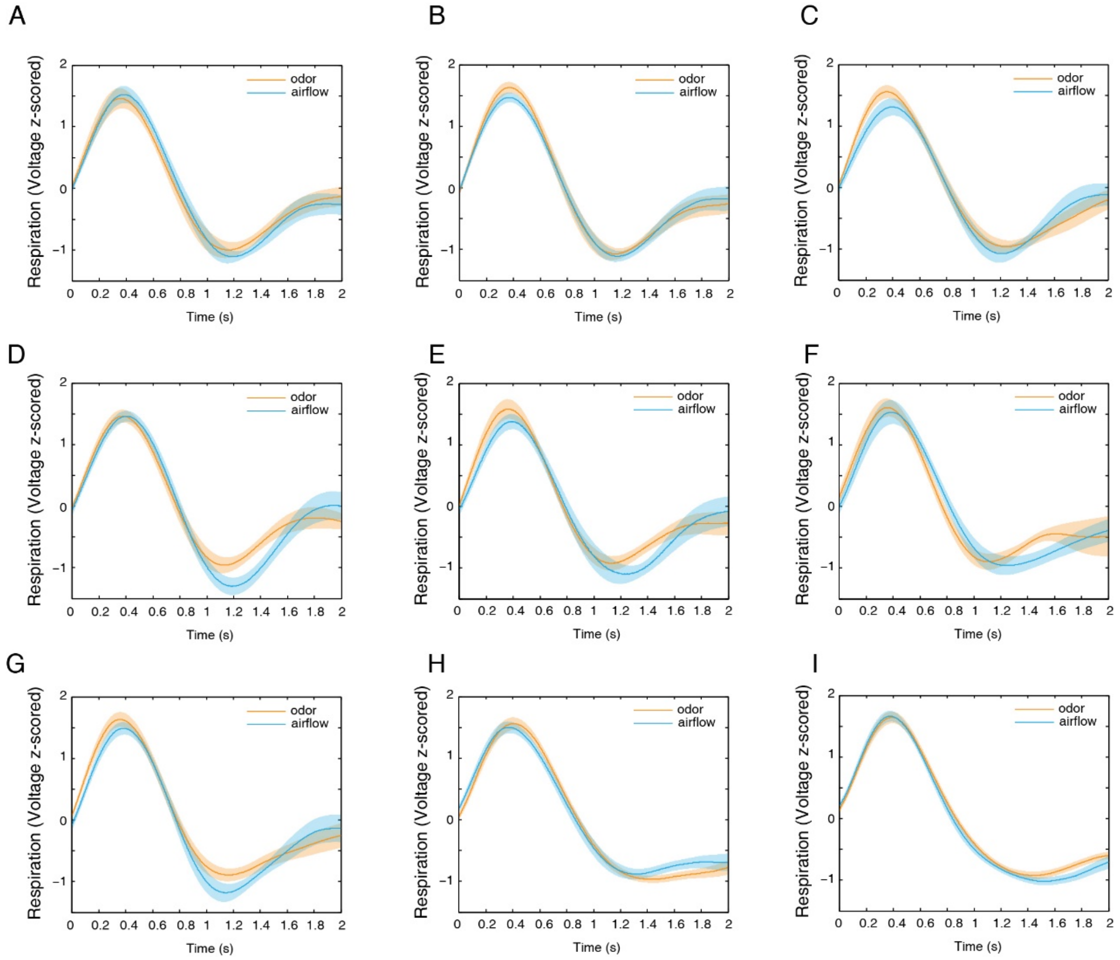


Figure S1: Participants' respiration state during odor discrimination task (odor) and airflow velocity testing (airflow). (A-G) Participants' respiration comparison in length difference of 5, 10, 15, 20, 25, 30 and 40cm in Exp. 2. The shadows are SEMs. (H, I) Participants' respiration comparison in temporal difference (H) and volume ratio difference (I) conditions in Exp. 3.
